# Compositional Canonical Correlation Analysis

**DOI:** 10.1101/144584

**Authors:** Jan Graffelman, Vera Pawlowsky-Glahn, Juan José Egozcue, Antonella Buccianti

**Affiliations:** Department of Statistics and Operations Research Universitat Politecnica de Catalunya Avinguda Diagonal 647, 08028 Barcelona, Spain. *email:*; Department of Biostatistics University of Washington UW Tower, 15th Floor, 4333 Brooklyn Avenue NE Seattle 98105 WA, USA; Department of Computer Science, Applied Mathematics, and Statistics Universitat de Girona Campus Montilivi, Edifici P4, E-17003 Girona, Spain. *email:*; Department of Civil and Environmental Engineering Universitat Politecnica de Catalunya Jordi Girona Salgado 1-3, Edifici C2, E-08034-Barcelona, Spain. *email:*; Department of Earth Sciences University of Florence Via G. La Pira 4, 50121 Firenze, Italy *email:*

**Keywords:** Canonical weights, canonical loadings, biplot, log-ratio transformation, generalized least squares, goodness-of-fit, biplot ray, biplot link, Moore-Penrose inverse

## Abstract

The study of the relationships between two compositions by means of canonical correlation analysis is addressed A coimnositional version of canonical correlation analysis is developed. and called CODA-CCO. We consider two approaches, using the centred log-ratio transformation and the calculation of all possible pairwise log-ratios within sets. The relationships between both approaches are pointed out, and their merits are discussed. The related covariance matrices are structurally singular, and this is efficiently dealt with by using generalized inverses. We develop compositional canonical biplots and detail their properties. The canonical biplots are shown to be powerful tools for discovering the most salient relationships between two compositions. Some guidelines for compositional canonical biplots construction are discussed. A geological data set with X-ray fluorescence spectrometry measurements on major oxides and trace elements is used to illustrate the proposed method. The relationships between an analysis based on centred log-ratios and on isometric log-ratios are also shown.

## 1 Introduction

Canonical correlation analysis (CCO) is an important classical multivariate method developed by Hotelling (1935; 1936) dedicated to the study of relationships between two sets of multiple variables, an *X*-set and a *Y*-set. Statistics courses on multivariate analysis usually cover the method, and textbooks in the field typically dedicate a chapter to the technique (Anderson 1984, Mardia et al. 1979, Johnson and Wichern 2002, Dillon and Goldstein 1984, Manly 1989). The monograph by Gittins (1985) is entirely dedicated to canonical analysis. Canonical correlation analysis offers a unifying theoretical framework, since several multivariate techniques are particular cases of it. CCO is a generalization of multiple regression with more than one response variable (Mardia et al., 1979; Gittins, 1985), relates to multivariate analysis of variance (MANOVA) and discriminant analysis when one of the two sets of variables consists of indicator variables (Gittins, 1985, Section 4.6), and is also intricately related to correspondence analysis (Greenacre, 1984, Section 4.4) when both the *X* variables and the *Y* variables consist of indicator variables. CCO has been greatly enhanced by the development of biplots that efficiently depict the correlation structure of the variables. The method provides a generalized least squares approximation to the between-set correlation matrix. Haber and Gabriel (1976), Ter Braak (1990) and Graffelman (2005) have shown that canonical correlation analysis allows the construction of a biplot of the between-set correlation matrix. The biplot greatly helps the interpretation of the output of a canonical correlation analysis.

Over the last decades, compositional data analysis (Aitchison, 1986) has experienced a strong development. Scientists have become increasingly aware of the fact that compositional data are special data and this has to be taken into account in any statistical analysis. Recent books by Pawlowsky-Glahn and Buccianti (2011), Van den Boogaart and Tolosana-Delgado (2013), and Pawlowsky-Glahn et al. (2015) show that the analysis of compositional data is an active field of research. Compositional data sets are inherently of multivariate nature. Log-ratio principal component analysis (Aitchison 1983) has become a standard multivariate technique in compositional data analysis, and is often one of the first tools used to explore a compositional data set. Specific compositional biplots (Aitchison & Greenacre 2002) have been proposed that allow efficient visualization of compositional data sets. Compositional data often goes together with other variables that can appear as predictors of the compositions, or that can appear as responses explained by compositions. In this paper we address the situation where there are two sets of variables which are both compositional in nature, and our goal is to study the relationships between the two sets by means of a canonical correlation analysis. Such data sets can arise in many settings. For example, in microbiological, nutritional and physiological studies it is of interest to study the relationships between food composition and body or microbial gut composition of the food consumers. In agricultural and ecological studies it is of interest to study the relationship between soil composition and plant composition. In geology, relationships between (sub)compositions with different types of compounds can be of interest. In biochemical studies often several kinds of components are measured such as proteins, carbohydrates, vitamins and fats. A compositional CCO allows one to study the relationships between the different kinds of compositions. The structure of this paper is as follows. In Section 2 we provide the theory for our compositional version of CCO, hereafter called CODA-CCO, and develop the corresponding compositional biplots. In Section 3 we illustrate our methodology with an artificial miniature example and with the analysis of a geological data set of major oxides and trace elements measured in European floodplain sediments. A discussion completes the paper.

## 2 Theory

In this section we establish our notation, briefly summarize classical CCO and then develop a compositional version of CCO.

### 2.1 Classical CCO

We consider one set containing p predictor variables (*X*-variables) and a second set containing *q* criterion variables (*Y*-variables). Both sets are assumed real, that is, the sample space is the ordinary Euclidean space. The *Y*-variables can be thought of as response variables, though not necessarily so, as the analysis treats *X* and *Y* in a symmetric fashion. The main aim of a CCO is to search for linear combinations **U** = **X**_c_**A** and **V** = **Y**_c_**B** of the column-mean centreed variables in **X**_c_ and **Y**_c_ that have maximal correlation. The coefficient matrices **A** and **B** are known as the *canonical weights* or the *canonical coefficients,* and the constructed linear combinations are known as the *canonical variables* (also termed *canonical variates* by some authors). The solution of a CCO is efficiently calculated by using the singular value decomposition (s.v.d.) of the transformed between-set covariance matrix. In particular, the canonical coefficients and correlations can be obtained by the s.v.d. of

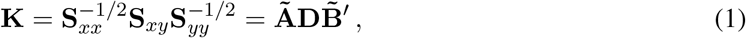

where **S**_xx_, **S**_yy_ and **S**_xy_ are the sample covariance matrices of the *X*-variables, the *Y*-variables, and the between-set covariances, respectively. Matrix *Ã* is a *p* × *r* orthonormal matrix of left singular vectors (**Ã′Ã** = **I**_r_) and matrix 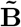 is a *q* × *r* orthonormal matrix of right singular vectors 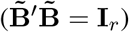. Diagonal matrix **D** is of rank *r* (*r* = min (*p*, *q*)) and contains the canonical correlations in non-increasing order of magnitude (Gittins, 1985, Section 2.3.2). The canonical coefficients are related to the left and right singular vectors by

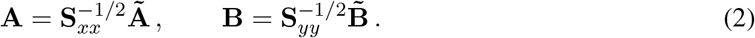

The canonical coefficients are normalized so that **A′S**_xx_**A** = **I**_r_ and **B′S**_yy_**B** = **I**_r_ and, consequently, the canonical variables are standardized variables,

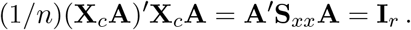

The singular value decomposition in (1) shows that we do a weighted least squares approximation of given rank to the between-set covariance matrix **S**_xy_. Row markers (**F**) and column markers (**G**) for the biplot can be obtained by:

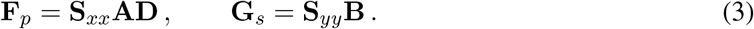

We use the subindices *p* and *s* to indicate “principal” and “standard” coordinates, respectively. This convenient terminology was proposed by Greenacre (1984) in the context of correspondence analysis, and was previously used in CCO by Graffelman (2005); it serves to distinguish the different biplot scalings. The principal coordinates are characterized by the presence of diagonal matrix **D** in the formula, whereas standard coordinates refer to coordinates without matrix **D** in their formula. An alternative scaling for the biplot is to have rows in standard coordinates, and columns in principal coordinates:

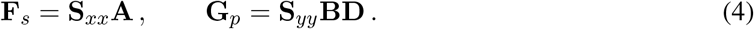

In CCO all these sets of coordinates for biplots can be interpreted as covariances. The principal coordinates **F**_*p*_ are cross covariances between *X*-variables and canonical *Y*-variables. The standard coordinates **G**_*s*_ are the covariances between canonical *Y*-variables and the original *Y*-variables. In the same manner, the standard coordinates **F**_*s*_ are intra-set covariances for the *X*-variables and the canonical *X*-variables, and the principal coordinates **G**_*p*_ are cross covariances between *Y*-variables and *X*-variables. This is shown by the following set of equations,

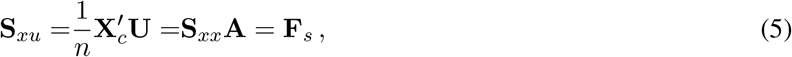

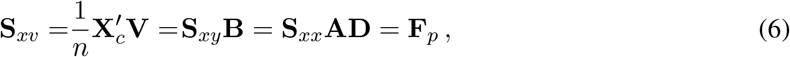

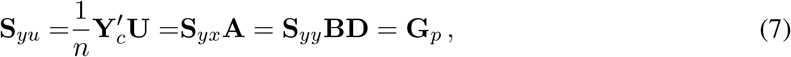

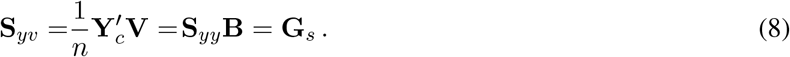

A biplot of the between-set covariance matrix **S**_xy_ can be obtained as **F**_*p*_**G**_*s*_**′** in Equation (3) or as **F**_*s*_**G**_*p*_**′** in Equation (4). Numerical output of a CCO typically also includes the *canonical loadings.* The canonical loadings are the correlations between the original variables and the canonical variables and can be used to interpret the canonical variables. In a correlation-based CCO the previous covariance expressions (Equations (5)–(8)) are in fact equal to the canonical loadings. If a covariance-based CCO is used, then the loadings are obtained by premultiplying the previous covariances with the inverse of a diagonal matrix containing the standard deviations (**D**_*sx*_, **D**_*sy*_), so that the loadings are obtained by:

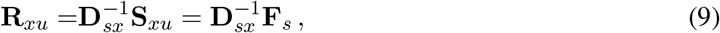

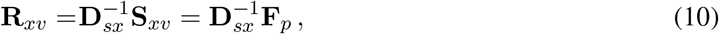

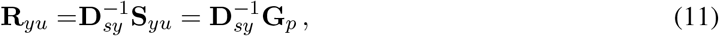

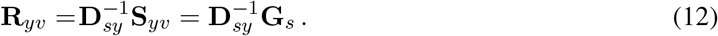

Note that, in order to obtain the loadings, post-multiplication by the inverse of the standard deviation of the canonical variables is not needed, as the latter are already standardized by virtue of the normalization constraints on the singular vectors in Equation (1). This shows that the correlation-based and covariance-based biplots are almost identical, and that the only difference is a rescaling of the variable vectors. In correlation-based CCO biplots all variable vectors will be within the unit circle. In covariance-based CCO biplots, variable vectors can be outside the unit circle. The angles between the variable vectors are the same in both types of analysis, and the goodness-of-fit of **S**_*xy*_ equals the goodness of fit of **R**_*xy*_.

We briefly summarize the main measures of goodness-of-fit in canonical analysis. The goodness-of-fit of the between-set covariance matrix in a k-dimensional biplot is given by

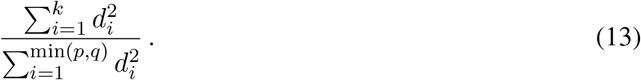

Matrix **X**_*c*_ is approximated by the inner products between the rows of **U** and the columns of **F**_*s*_. If the *X*-set is the smaller set (*p* ≤ *q*) then, in the full space of the solution, **X**_*c*_ is perfectly recovered, because

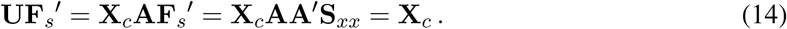

In a *k*-dimensional biplot **X**_*c*_ is approximated by 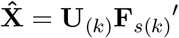. The total variance of the **X** variables accounted for by a given number of canonical **U** variables, called the *adequacy coefficient* (Thompson, 1984), is

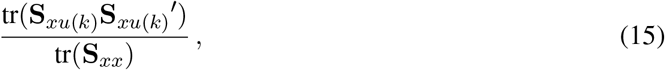

where **S**_*xu*(*k*)_ refers to the covariance matrix between *X* variables and the first *k* canonical *X* variables. Note that the adequacy coefficient is not scale-invariant under standardization of the original variables. With standardized variables, the adequacy coefficients are obtained by changing all covariances in Equation (15) by correlations, and this reduces to 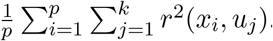. The latter is also the average of the coefficients of determination (*R*^2^) obtained by regressing all *X* variables onto *k* canonical variables. Likewise, the inner products of *U* with the *Y*-variables in principal scaling approximate the *Y*-measurements in the full space, and we have

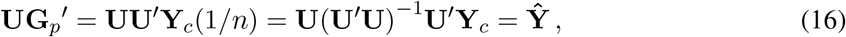

which can be interpreted as the fitted values obtained in a regression of **Y**_*c*_ onto the canonical *X*-variables. In general, it will not be possible to exactly recover the measurements of the variables in principal scaling, even if we use the full space of the solution. The amount of explained variation of the *Y*-variables in a *k*-dimensional solution, known as the *redundancy coefficient* (Stewart and Love, 1968), is

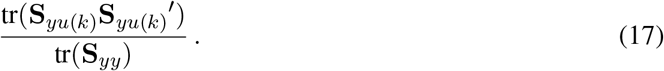

The redundancy coefficients are neither scale-invariant under standardization of the original variables. With standardized variables, the redundancy coefficients are obtained by changing the covariances in Equation (17) by correlations, and reduce to 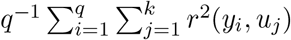. Analogous adequacy and redundancy coefficients can be calculated for the canonical *Y* variables.

In conclusion, classical CCO basically provides a biplot of the between-set covariance or correlation structure, in which the original observations are absent. Classical biplots made by principal component analysis (Gabriel, 1971) provide more information, since they do not only represent the variables, but also the original samples. In previous work, Graffelman (2005) has shown that it is possible to represent the original samples in the CCO biplot by using regression results for the representation of supplementary information (Graffelman and Aluja-Banet, 2003). If samples are fitted to the biplot by generalized least squares, it is particularly simple to represent them in the biplot: the **F**_*s*_**G**_*p*_′ biplot should be overplotted with the canonical *X* variables, and the **F**_*p*_**G**_*s*_′ biplot should be overplotted with the canonical *Y* variables. These results are of particular relevance for a compositional version of CCO, as they will allow the representation of the original compositions in the CCO biplot (See Section 2.2). The corresponding plots could be termed *triplots* because they represent three entities: *X* variables, *Y* variables and data points. The term triplot stems from ecological multivariate analysis, as triplots are commonly made in canonical correspondence analysis (Ter Braak, 1986; Ter Braak and Smilauer, 2002) and redundancy analysis (Ter Braak and Looman, 1994).

We finish this section with a few remarks on the scaling of the original data matrix, as this is also relevant for the compositional analysis that is to follow. One can decide to perform CCO using covariance matrices (as outlined above), or using correlation matrices. A correlation based analysis is possible by simply standardizing the data matrices prior to the analysis, e.g. dividing the columns of **X** and **Y** by their respective standard deviations. CCO is, to a large extent, invariant to such standardization. Canonical correlations, canonical variables, and canonical loadings will all be the same in a covariance-based and a correlation-based analysis. In this sense CCO differs from principal component analysis (PCA), being it well known that a PCA of the centreed data matrix is different from the PCA of the standardized data matrix, giving rise to two “variants” of PCA. The main difference between a covariance based CCO and a correlation based CCO concerns the biplot: the first produces a biplot of the between-set covariance matrix, whereas the latter produces a biplot of the between-set correlation matrix. The goodness-of-fit of these matrices will be the same in both approaches. Finally, the goodness-of-fit of the original data matrices, as expressed by the adequacy and redundancy coefficients, differ in a covariance-based and correlation-based analysis as explained above.

### 2.2 Compositional CCO

In the development in the previous section, **X** and **Y** typically stand for matrices of quantitative real variables. We now consider **X** and **Y** to be matrices with n compositions in their rows, and having *D_x_* and *D_y_* parts (columns) respectively. Recall that compositional data can be defined as strictly positive vectors for which the information of interest is in the ratios between components. There are several ways to perform a CODA-CCO, depending on how the compositions are transformed. One can use the additive, the centred or the isometric log-ratio transformation, or one can also use the matrices with all pairwise log-ratios of the *X*-set and the *Y*-set. The different approaches are largely equivalent, though the biplots obtained will be different. We develop two approaches to CODA-CCO in the corresponding subsections below, using the canonical analysis of the clr transformed compositions (2.2.1), and, largely equivalently, the canonical analysis of all pairwise log-ratios of the *X*-set and the *Y*-set (2.2.2). Both these transformations lead to a visualization of the pairwise log-ratios which form the most simple representation of the data, and from which more complex ratios can be build. The clr-based approach is also the usual approach take in log-ratio principal component analysis (Pawlowsky-Glahn et al., 2015; Aitchison and Greenacre, 2002). Some invariance properties for the isometric log-ratio transformation are derived in Appendix A.

#### 2.2.1 The centred log-ratio (clr) approach

We consider the centred log-ratio transformation (clr) of a composition x given by

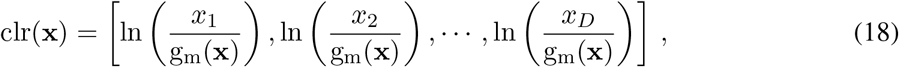

where g_m_(x) is the geometric mean of the components of the composition x. Let **X**_*l*_ be the log transformed compositions, that is **X**_*l*_ = ln (**X**) with the natural logarithmic transformation applied elementwise. The clr transformed data can be obtained by just centring the rows of this matrix, using the centring matrix 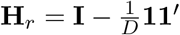, with *D* equal to *D_x_* or *D_y_* as corresponds. Then

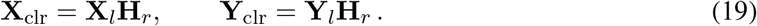

These clr transformed data matrices have the same dimensions as **X** and **Y**. The columns of **X**_clr_ and **Y**_clr_ are subject to a zero sum constraint because **H**_r_**1** = 0. The column rank of these matrices is, in the absence of additional linear constraints, equal to *D_x_* – 1 and *D_y_* – 1, respectively. We now column-centre the clr transformed data, producing data matrices that have column means that are zero,

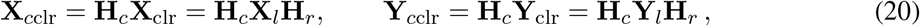

where **H**_c_ is the idempotent centring matrix **H**_*c*_ = **I** – (1/*n*)**11**′. Thus, **X**_cclr_ and **Y**_cclr_ have zero row means due to the subtraction of the geometric means, and zero column means due to centring operation **H**_c_. We propose to use **X**_*c*clr_ and **Y**_*c*clr_ as the input matrices for a classical covariance-based CCO described in Section 2.1. Due to the zero row sum constraint, the covariance matrices of **X**_*c*clr_ and **Y**_*c*clr_ are singular. In CCO the covariance (or correlation) matrices of the *X* and *Y* variables are inverted. In order to be able to deal with the structural singularity due to the compositional nature of the data, we use a generalized inverse, the Moore-Penrose inverse (Searle, 1982), in order to be able to proceed with the analysis. In CCO with non-singular covariance matrices, the inverse of the square roots of the covariance matrices are needed (Equation 1) and these can be obtained from the spectral decomposition of the covariance matrices, in particular

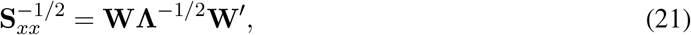

where **W** and **Λ** contain eigenvectors and eigenvalues obtained in the spectral decomposition of **S**_*xx*_ = **WΛW**′. Under singularity of **S**_*xx*_, the Moore-Penrose inverse denoted by 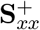 is obtained by 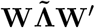, with 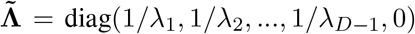, which satisfies the four Moore-Penrose conditions. Compositional canonical correlation analysis (CODA-CCO) can then be carried out using the singular value decomposition

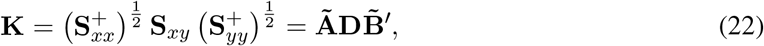

Due to the compositional nature of the data, the number of dimensions in the solution, the rank of **D**, is now given by *r* = min (*D_x_* – 1, *D_y_* – 1). The canonical coefficients are now obtained as

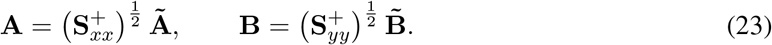

The biplot coordinates and the canonical loadings of a CODA-CCO are now obtained by the same expressions given for the classical analysis in Equations (3) and (4) and (5) through (8). Note that the between-set covariance matrix of the clr coordinates **S**_*xy*_ has dimension *D_x_* × *D_y_*, but that the generalized inverses have at most rank *D_x_* – 1 and *D_y_ – 1*, respectively. Consequently, matrix **K** is not full rank, but has at most rank min(*D_x_* – 1, *D_y_* – 1). We note that computer programs typically produce an s.v.d. where **D** has dimensions (*r* + 1) × (*r* + 1), implying that **D** has a trailing zero on the diagonal which is consequence of the singularity of the covariance matrices of the clr transformed data. If the s.v.d. in (22) is conceived that way, the corresponding normalization of the canonical coefficients is affected, and one has that **A′S**_*xx*_**A** = **Ĩ** and **B′S**_*yy*_**B** = **Ĩ**, where **Ĩ** is a diagonal matrix with *r* ones and one trailing zero on its diagonal. In the remainder, we conceive **D** of dimension *r* × *r*, without trailing zero, such that the canonical coefficient matrices have no trailing column of zeros and can be considered to be full column rank, and satisfy the usual normalizations **A′S**_*xx*_**A** = **I**_r_ and **B′S**_*yy*_**B** = **I**_*r*_. Note that the columns of the matrices of canonical coefficients sum to zero. A justification for this is given in appendix A. We complete this section enumerating some properties of the compositional canonical biplots obtained. Most of these properties are relatively straightforward extensions of the previous work on compositional biplots (Aitchison and Greenacre, 2002) and canonical biplots (Haber and Gabriel, 1976; Ter Braak, 1990; Graffelman, 2005). For a treatment of compositional biplots, see also Section 5.4 of Pawlowsky et al. (2015).

1. Biplot origin. The origin of the biplot represents the vector of geometric means of the n compositions. In **F**_*s*_**G**_*p*_′ scaling the origin corresponds to the geometric mean vector of the *X* compositions, whereas in **F**_*p*_**G**_*s*_′ scaling, the origin corresponds to the geometric mean vector of the *Y* compositions. This can be seen from equations **U** = **X**_*c*clr_**A** and **V** = **Y**_*c*clr_**B**. If the double centring operation is applied to the vector of geometric means, a zero vector is obtained, and consequently the values of the canonical variables are zero. At the same time, the origin of the biplot is also the point from which the biplot vectors representing the clr components emanate.
2. Biplot vector (ray) length. Due to the symmetric nature of CCO, we can assume *D_x_* ≤ *D_y_* without loss of generality. The length of variable vectors plotted in standard coordinates is, for the smallest composition (the one with fewer parts), in the full space of the solution, equal to the standard deviation of the corresponding clr transformed part. This follows from

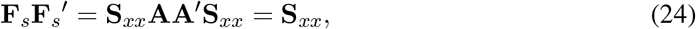

where the last equality follows from the fact that **AA**′ is the Moore-Penrose inverse of **S**_*xx*_. If a two-dimensional biplot is used as an approximation of the data set, ray length will underestimate the observed sample standard deviation. It also follows that the length of a biplot vector can never exceed the sample standard deviation of the corresponding clr component. For the larger composition (the one with more parts), we have, in the full r dimensional space

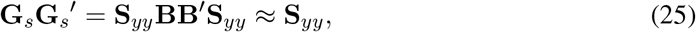

where the LHS has rank *r*, but **S**_*yy*_ has rank *D*_*y*_ – 1 ≥ *r*. Thus, for the larger composition, the length of the rays will be smaller than the standard deviation of the corresponding clr transformed part. Finally, biplot rays of parts that are plotted in principal coordinates are shrunk with respect to the standard coordinates due to the postmultiplication by the canonical correlations (see Equations (3) and (4)) and will always fall short of the observed sample standard deviation, and give a worse approximation to it compared with the standard coordinates. This is consistent with previous work (Graffelman, 2005), where it was shown that the within-set covariance matrices are better approximated with biplot vectors in standard coordinates.
3. Inner products between biplot vectors. It follows from Equation (24) that the inner product between two biplot vectors of the *same* set (again in the full space, using standard coordinates, and correspondingly the set with the smaller composition) equals the covariance of the corresponding clr components. Inner products of biplot vectors *between* subsets (one set in standard and the other set in principal coordinates) approximate the between-set covariance matrix of clr transformed parts. This is justified by

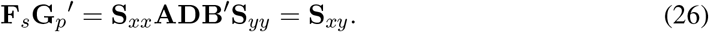 This approximation is optimal in the generalized least squares sense as guaranteed by the s.v.d in Equation (22), and it is the same in both biplot scalings, and in fact the focus of the analysis.
4. Cosine of angle between two biplot vectors. The cosine of the angle between the two vectors *within sets* (again referring to the standard coordinates of the smaller composition) equals the sample correlation of the clr components in the full space. In a two-dimensional subspace this will be “approximately so”, being it unknown if the approximation is optimal in some sense. Cosines of angles of biplot vectors *between* subsets will exaggerate the correlations between transformed clr components of the two subsets, even in the full space of the solution. This is because the ray lengths of the larger composition underestimate the standard deviation of the corresponding part (see the previous point 2). Importantly, the approximation to the correlations offered by using cosines depends on the biplot scaling. It is not the same in the **F**_*s*_**G**_*p*_′ and the **F**_*p*_**G**_*s*_′ scaling. This is because the length of the biplot vectors in the rows of **G**_p_ and **F**_p_ fall short of the corresponding standard deviation to a different extent.
5. Link length. A biplot link is the difference vector of two biplot rays. In CODA biplot interpretation, the links are very important because they represent the log-ratio of the connected parts. For the composition that is represented in standard coordinates, the length of a link in the full space of the solution equals the standard deviation of the corresponding log-ratio. Let f_i_ and f_j_ represent the rays of parts *i* and *j* respectively (rows of **F**_s_). The squared length of their link is given by

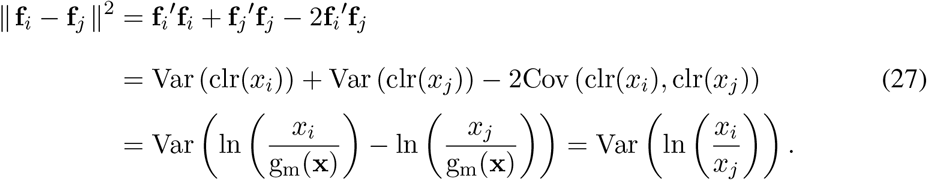 Under the considered scaling, the links of the larger composition will necessarily be represented in principal coordinates. Let g_i_ and g_j_ represent the rays of parts *i* and *j* respectively (rows of **G**_*p*_). The squared length of their link is given by

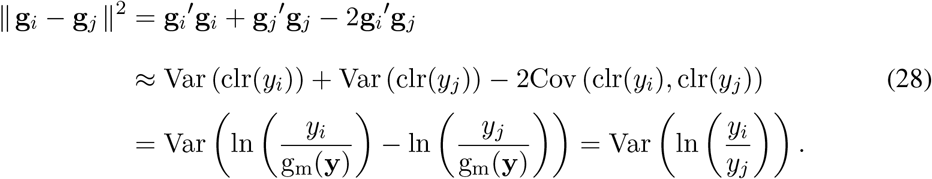 This shows there is no corresponding full space result for the length of the links in principal coordinates (note the use of ≈ in the last equation). As argued above, the terms g_i_′g_i_ and g_j_′g_j_ underestimate the corresponding standard deviation, even in the full space. The principal links will equal the corresponding standard deviations in the full space only in the case of equally sized compositions (*p* = *q*) and all canonical correlations equal to 1.
6. Inner products between links. Since the focus of the analysis is on relationships between the log-ratios of the two sets, inner products and angles between *X* and *Y* links are of interest. Links are vectors of differences, and the inner product between two links corresponding to the log-ratios ln (*x_i_/x_j_*) and ln (*y_r_/y_s_*) is, in full space, the covariance between the two corresponding log-ratios because

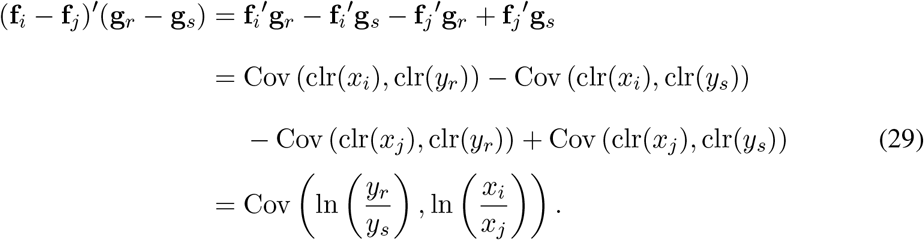 This equation is exact in the full space and has interesting implications. Since all four clr covariances are optimally approximated in the analysis, the implication is that the covariances between log-ratios of the *X* set and the *Y* set are also optimally approximated. An alternative way to construct a CODA-CCO biplot is then to depict only links as arrows emanating from the origin and leave the clr components out of the biplot (e.g. see Figures 2A and 3A in the Example section, where the links in 2A are identified as the rays in 3A), this gives precisely the CODA-CCO biplot obtained in the pairwise log-ratio approach (See subsection 2.2.2).
7. Cosines of angles between links. Equations (27), (28) and (29) show that, in the full space, cosines of angles between links are “close to” the correlations of the corresponding log-ratios. However, because of the aforementioned inexact nature of Equation (28), cosines of angles will not equal sample correlations between log-ratios exactly.

Up to this point, CODA-CCO has been developed using a covariance-based approach, mainly because all clr transformed parts have the same log-ratio scale. This implies that inner products in the CODA-CCO biplots (Equations (24) through (26)) represent covariances between clr transformed parts as well. Covariances are only indicative of the nature of the relationship (direct or indirect) but not about the strength of the observed relationship. For the latter purpose, correlations are far more useful. From the foregoing it is clear that in CODA-CCO the approximation of the correlations by cosines is problematic for two reasons: first, for being inexact in the full space (when the larger composition is considered, or when principal coordinates are involved), and second, for having no justification that approximations in low-dimensional biplots are optimal. In order to avoid these problems, one might therefore consider to standardize the clr transformed data, such that the scalar products in Equations (24) through (26) will approximate the correlations. This however, yields a biplot that approximates correlations between clr transformed parts, which do not seem particularly interesting. Note that the covariance on the right hand side of Equation (29) is *not* converted into a correlation by standardizing the clr data. Potentially more interesting biplots, tightly related to the clr approach exposed here, are obtained in the pairwise log-ratio approach in the next section.

#### 2.2.2 The pairwise log-ratio (plr) approach

An alternative approach to CODA-CCO is to use the pairwise log-ratios (plr for short) of the *X*-set and the *Y*-set, and to submit these to a canonical analysis. First, we define two matrices *X_p1r_* and *Y_plr_* with all possible log-ratios for the *X* and *Y* set respectively, having dimensions 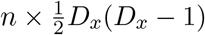 and 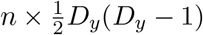 respectively. We column-centre these matrices to obtain

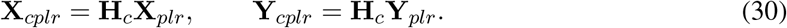

CODA-CCO is now performed by the s.v.d. of the transformed between-set covariance matrices of **X**_*cp1r*_ and **Y**_*cpir*_, that is, by applying Equation (22) to the covariance matrices of the newly defined data matrices. Because of the structural singularity of **S**_*xx*_ and **S**_*yy*_, again the Moore-Penrose inverse of the latter two is used. It is immediately clear that the clr-approach and plr-approach are “equivalent” to a large extent. Any pairwise log-ratio is a linear combination of the clr transformed parts because

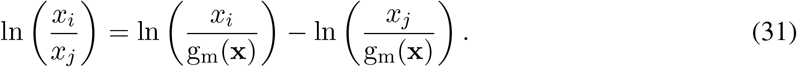

It therefore follows that **X**_*plr*_ and **Y**_*plr*_ have the same rank as **X**_clr_ and **Y**_clr_ respectively, and the number of dimensions with non-zero singular values is the same in both analysis. Moreover, Equation (29) already showed that the covariances of the plr data are linear combinations of the covariances of the clr data. Canonical correlation analysis is known to be invariant under linear transformations of the data. It is thus clear that the canonical correlations and the canonical variables obtained are the same in both types of analysis. The canonical coefficients are however, not invariant, and this implies, by virtue of Equations (3) and (4), that the biplot is affected. In the plr approach, biplots will generally be crowded with more rays, 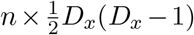 and 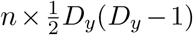, respectively, for each set. These biplot vectors now *directly* represent the pairwise log-ratios. In the plr approach, biplots properties are straightforward to infer using the results in subsection 2.2.1. We express these therefore more concisely, but emphasize some novelties.

1. Biplot origin. The origin of the biplot now represents the mean of each pairwise log-ratio, both for the pairwise log-ratios of the *X* set and the *Y* set.
2. Biplot vector (ray) length. The length of a variable vector plotted in standard coordinates is, for the smallest composition, in the full space of the solution, according to Equation (24) now equal to the standard deviation of the corresponding log-ratio. Correspondingly, ray lengths in standard coordinates for the larger composition will underestimate the standard deviation of the corresponding log-ratio. Also correspondingly, biplot rays of parts plotted in principal coordinates give poorer approximation of the corresponding standard deviations of the the log-ratios.
3. Inner products between biplot vectors. Equation (24) now shows, with again the same conditions (full space, standard coordinates, the smaller composition), that the inner product between two biplot vectors of the *same* set equals the covariance of the corresponding log-ratios. Inner products of biplot vectors *between* subsets approximate the between-set covariance matrix of log-ratios, the latter being optimal in the generalized least squares sense.
4. Cosine of angle between two biplot vectors. The cosine of the angle between the two vectors *within sets* (standard coordinates, the smaller composition, full space) equals the sample correlation between two log-ratios. Cosines of angles of biplot vectors *between* subsets exaggerate the correlations between the log-ratios of the two subsets and depend on the biplots scaling for reasons previously described.
5. Links. A biplot link now becomes the difference vector of two log-ratios. If the two log-ratios share a part, having it both in the numerator, or both in the denominator, the link is another log-ratio because

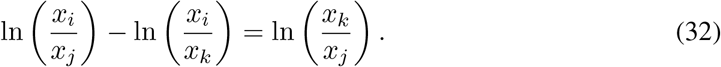 Representing this link is superfluous, as the biplot already shows all pairwise log-ratios as vectors emanating from the origin. If the two log-ratios don’t share parts, we have

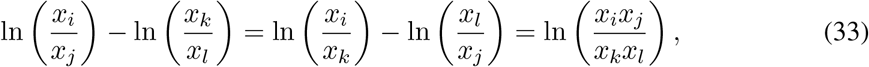

showing that the biplot will have identical, duplicated links, to be interpreted as “differences in log-ratios”. If the two log-ratios share a part, one having it in the numerator and one having it in the denominator, we have

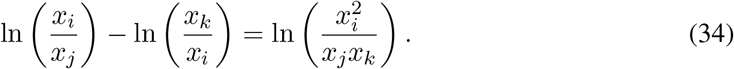 Equation (32) is a simple log-ratio, whereas Equations (33) and (34) are examples of *balances* (Pawlowsky-Glahn et al., 2015). Balances can be very useful and can have substantive interpretation depending on the context of the data being analysed. At this point we refrain from developing inner products and cosines for links in the pairwise approach, and will focus mainly on the rays (pairwise log-ratios) for interpretation.

We argued above that in the clr approach standardization of the data did not seem very useful. In the plr approach, standardization can be highly useful, and it is probably often to be recommended. The reason is that standardization of the pairwise log-ratios now converts Equations (24), (25) and (26) into correlation matrices. In particular, Equation (26) implies the biplot can now efficiently visualize the correlation structure of the pairwise log-ratios, and that optimal low-dimensional approximations to this correlation structure can be obtained. This was not possible in the clr approach given in Section 2.2.1.

## 3 Examples

In this section we present two examples of a compositional canonical correlation analysis. The first example concerns two synthetic 3-part compositions registered for the same set of subjects. The advantage of this example is that the between-set covariance matrix is of rank two, and that everything can be represented without error in two-dimensional space. The second example is geological and concerns the chemical composition (major oxides and trace elements) of European floodplain sediments.

### 3.1 Two sets of compositions of three parts

We show 100 observations on two 3-part compositions, *x* and y, in the ternary diagrams in Figure 1. The ternary diagram of the *X*-set reveals a clear pattern, having an approximately constant *x*_1_/*x*_2_ ratio, whereas the *Y*-set shows, at first sight, no clear structure. These ternary plots only reveal marginal information on the *X* and *Y* compositions, and are not informative about the relationships between the *X*-set and the *Y*-set. We calculate the centred log-ratio transformation of compositions *x* and *y* separately, and perform the clr-based compositional canonical correlation analysis developed in the previous section. Table 1 shows the classical numerical output of a CCO analysis. Initially, we use a covariance-based analysis, because all variables are in a commensurable log-ratio scale.

**Figure 1:**
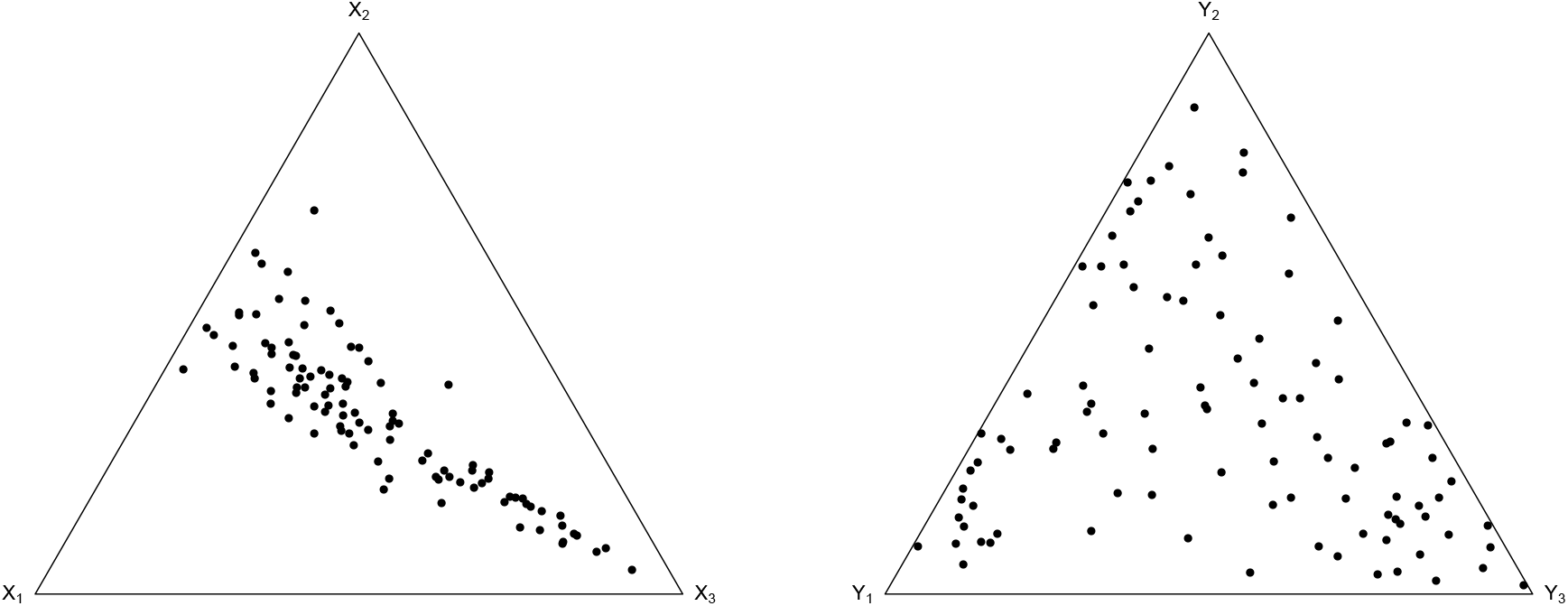
Ternary diagrams of two compositions, x and y, of three parts.

**Table 1:** Canonical correlations (*r*_1_, *r*_2_), canonical weights, canonical loadings (between parentheses), adequacy coefficients (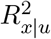, and cumulative 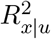) and redundancy coefficients (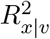, and cumulative 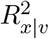) obtained in a CODA-CCO of two sets of clr transformed compositions of three parts.

| | $r_1 = 0.944$ | | $r_2 = 0.129$ | |
| --- | --- | --- | --- | --- |
| | $U_1$ | | $U_2$ | |
| $\text{clr}(x_1)$ | 0.001 | (-0.886) | 3.847 | (0.464) |
| $\text{clr}(x_2)$ | -0.799 | (-0.983) | -3.447 | (-0.184) |
| $\text{clr}(x_3)$ | 0.798 | (0.994) | -0.401 | (-0.109) |
| $R_{x\vert u}^2$ | 0.913 | | 0.087 | |
| $R_{x\vert u}^2$ | 0.913 | | 1.000 | |
| $R_{x\vert v}^2$ | 0.813 | | 0.001 | |
| $R_{x\vert v}^2$ | 0.813 | | 0.815 | |
| | $V_1$ | | $V_2$ | |
| $\text{clr}(y_1)$ | 0.762 | (0.852) | -0.050 | (-0.523) |
| $\text{clr}(y_2)$ | -0.717 | (-0.610) | -0.521 | (-0.793) |
| $\text{clr}(y_3)$ | -0.046 | (-0.303) | 0.572 | (0.953) |
| $R_{y\vert v}^2$ | 0.397 | | 0.603 | |
| $R_{y\vert v}^2$ | 0.397 | | 1.000 | |
| $R_{y\vert u}^2$ | 0.353 | | 0.010 | |
| $R_{y\vert u}^2$ | 0.353 | | 0.363 | |

Table 1 shows that the first canonical correlation is very high, 0.94, implying that the two variable sets share a large part of their variation. All of the variance of clr(x) and clr(y) is accounted for by the two canonical variables, as expected. The goodness-of-fit of the between-set covariance matrix **S**_*xy*_ is also 100 percent, as predicted. Considering only one dimension, it is 0.944^2^/(0.994^2^ + 0.129^2^) = 0.982. This suggests there is only one important dimension. The cumulative adequacy coefficients 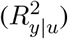 show that a two-dimensional **F**_*s*_**G**_*p*_′ biplot explains 100% of the variance of the clr transformed *X* parts, and 36.3% of the variance of the clr transformed *Y* parts. Most of the variance of the clr transformed parts is accounted for by the first dimension of the analysis. This dimension accounts for 91.3% of the variance in the transformed *X* parts, and for 35.3% of the variance in the transformed *Y* parts. The first canonical variate *U*_1_ correlates strongly with all *X* parts, and *V*_1_ correlates strongly with *y*_1_ and *y*_2_. The second canonical correlation is small, and non-significant in a permutation test (see below). Log-ratio CODA-CCO biplots are shown in various scalings in Figure 2. Biplots have been overplotted with the canonical variables (multiplied by a single convenient scaling factor, using the rows of matrix **U** in Figures 2A and 2C, and the rows of matrix *V* in Figures 2B and 2D) in order to represent the original compositions in the biplot. The variable labels *X_i_*, *Y_j_* in the plot actually represent the clr transformed parts. A link between rays *i* and *j within* a subset represents the corresponding log-ratio ln (*x_i_/x_j_*). The key point of these biplots is to look for *parallel links of each subset that run parallel to a canonical variable* with a high correlation. The canonical variables “channel” the correlation structure of the variables and represent the most correlated feature of the data. Figure 2A shows parallel links between (clr(*x*_1_), clr(*x*_2_)) and (clr(*y*_2_), clr(*y*_3_)), implying that the log-ratios ln(*x*_1_/*x*_2_) and ln(*y*_2_/*y*_3_) are correlated. However, the corresponding link is not parallel to the first canonical variable, and these log-ratios have only weak correlation. Moreover, Figures 2B, 2C and 2D do not show this parallelism, suggesting that it is accidental. More interestingly, Figure 2A also shows long parallel links through (clr(*_x_*_2_), clr(*_x_*_3_)) and through (clr(*_y_*_1_), clr(*_y_*_2_)) that run parallel to the first canonical variate, suggesting that the log-ratios ln(x_2_/x_3_) and ln(y_1_/y_2_) are highly correlated. These interpretations are confirmed by the sample correlations between these log-ratios; _r_(ln(x_1_/x_2_), ln(y_2_/y_3_)) = –0.14 and _r_(ln(x_2_/x_3_), ln(y_1_/y_2_)) = —0.94. Correlations inferred from the biplot can be corroborated by making a scatterplot matrix of all possible log-ratios, as is shown in supplementary Figure S1. An additional approximately parallel pair of links with some inclination is observed in Figure 2A between (clr(*x*_1_), clr(*x*_3_)) and (clr(*y*_1_), clr(*y*_3_)). The corresponding log-ratios have a correlation of −0.56. The presence of the samples in the biplot aids interpretation and illustrates the observed correlations: the compositions projecting high onto the link through (clr(*x*_2_), clr(*x*_3_)) also project high onto the link through (clr(*y*_1_), clr(*y*_2_)) and so confirm the correlated nature of the corresponding log-ratios.

**Figure 2:**
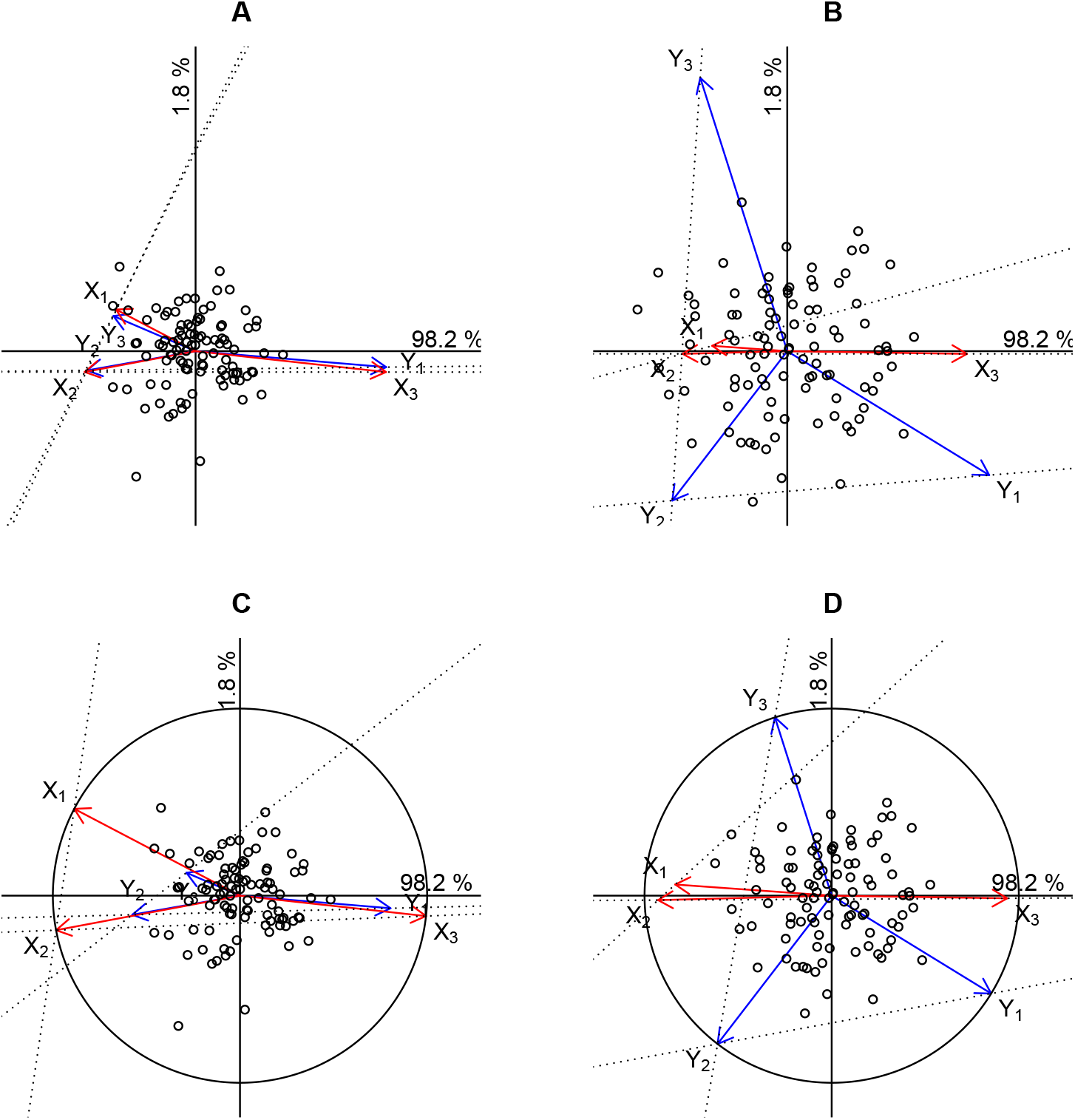
CODA-CCO clr biplots of two three-part compositions using different scalings. Rays represent clr-transformed parts. Links (clr(*x*_1_), clr(*x*_2_)), (clr(*x*_2_), clr(*x*_3_)), (clr(*y*_1_), clr(*y*_2_)), (clr(*y*_2_), clr(*y*_3_)) are indicated by dotted lines. Panels A (FsGp^’^ scaling) and B (FpGs^’^ scaling) are biplots made with a covariance-based analysis. Panels C (**F**_*s*_**G**_*p*_′ scaling) and D (**F**_*p*_**G**_*s*_′ scaling), with unit circle, are biplots made with a correlation-based analysis.

An alternative biplot for the same data, using the **F**_*p*_**G**_*s*_′ scaling from Equation 3, is shown in Figure 2B. This biplot explains 81.5% of the variance of the clr transformed *X* parts, and 100% of the variance of the clr transformed *Y* parts. The goodness-of-fit of the between-set covariance matrix is the same as in Figure 2A (100%). However, the biplot in Figure 2B seems to be the more interesting option if the original compositions are added to the biplot, because overall it accounts for more variability of the clr transformed data. Note that the links corresponding to the log-ratios ln(*x*_1_/*x*_2_) and ln(*y*_2_/*y*_3_) are now not far from orthogonal, whereas in Figure 2A they were virtually parallel. This shows that one needs to be cautious when interpreting the biplot, and that parallelism of links does not necessarily imply strong correlation of the corresponding log-ratios. Also note that the links corresponding to the log-ratios ln(*x*_2_/*x*_3_) and ln(*y*_1_/*y*_2_) are close to parallel in the direction of the first canonical variate, and that this is observed in *both* biplot 2A and 2B. This is the most salient relationship between the two compositions. Figure 2B also shows almost horizontal parallel links through (clr(*x*_1_), clr(*x*_3_)) and (clr(*y*_1_), clr(*y*_2_)), and more clearly reveals the correlation between the corresponding log-ratios. Figures 2C and 2D show CODA-CCO biplots of the same data, but with the clr-data standardized prior to the canonical analysis. In these plots, scalar products between the sets correspond to correlations between the clr components of each set. These plots resemble Figures 2A and 2B, but with rescaled rays. This is precisely what is expected as a consequence of the invariance of CCO under linear transformation. Note that the goodness-of-fit of the between-set covariance matrix and the between-set correlation matrix is the same as expected. However, plots 2C and 2D do add value to the previous graphs in two ways: firstly, due to the presence of the unit circle it is possible to infer that the clr transformed *X* components are perfectly represented in Figure 2C and the *Y*-parts in Figure 2D. Secondly, Figures 2C and 2D provide optimal approximations of the between-set correlation structure of the clr transformed components, whereas Figures 2A and 2B do not. Because of the small size of the miniature example, and because *D_x_* = *D_y_* = 3, cosines of angles in Figures 2A and 2B do coincide with the between-set sample correlations, but for larger compositions with *D_x_* = *D_y_* this will generally not be the case.

CODA-CCO biplots that are based on the analysis of pairwise log-ratios are shown in Figure 3. Now, each biplot vector represents a log-ratio. Due to the aforementioned invariance, goodness-of-fit of the covariance and correlation matrices of the log-ratios is the same as in the previous clr-based approach. Note that in Figures 3A and 3B, each biplot vector equals the sum or difference of the other two vectors of its set, which is a consequence of Equation (32).

**Figure 3:**
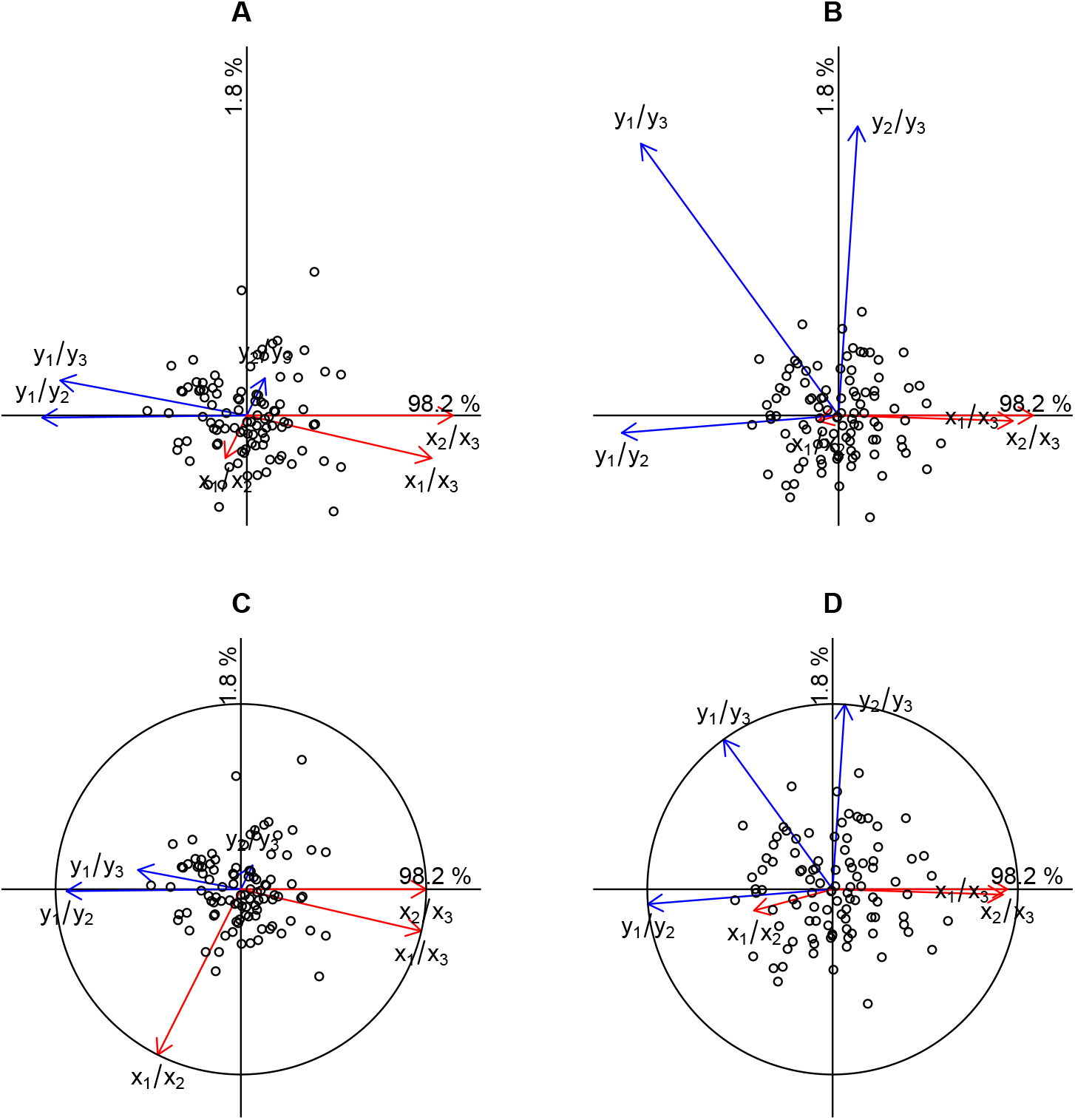
CODA-CCO biplots using pairwise log-ratios of two three-part compositions using different scalings. Rays represent log-ratios. Panels A (**F**_*s*_**G**_*p*_′ scaling) and B (**F**_*p*_**G**_*s*_′ scaling) are biplots made with a covariance-based analysis. Panels C (**F**_*s*_**G**_*p*_′ scaling) and D (**F**_*p*_**G**_*s*_′ scaling), with unit circle, are biplots made with a correlation-based analysis.

In all biplots in Figure 3 the log-ratios ln (*x*_2_/*x*_3_) and ln (*y*_1_/*y*_2_) virtually coincide with the first canonical variate. Indeed, the first canonical variate can be interpreted as the difference between these two log-ratios, and confirms this is the most correlated aspect of the data. Results obtained with standardization of log-ratios shown in Figures 3C and 3D do leave angles between vectors unaltered, but this is only because Figure 3 represents full space results with *D_x_* = *D_y_*. Between-set inner products in 3C and 3D are now correlations and identify ln (*y*_2_/*y*_3_) as uncorrelated with all log-ratios except ln (*y*_1_/*y*_3_).

### 3.2 Geochemical data: the composition of European floodplain sediments

The analysed data base is given by the chemical composition of floodplain sediments and is drawn from the FOREGS Geochemical Baseline Mapping Program initiated in 1998 to provide high quality environmental geochemical baseline data in Europe (http://weppi.gtk.fi/publ/foregsatlas/). The data set consists of 747 samples, stratified by European country. The concentrations, expressed as weight % for 10 major oxides (SiO_2_, Al_2_O_3_, Na_2_O, MgO, P_2_O_5_, K_2_O, CaO, TiO_2_, MnO, Fe_2_O_3_) and in ppm for 18 trace elements (V, Cr, Co, Ni, Cu, Zn, Ga, As, Rb, Sr, Zr, Nb, Sn, Cs, Ba, Pb, Th, U), were determined by X-Ray Fluorescence (XRF) spectrometry. XRF spectrometry is one of the most widely used and versatile of all instrumental analytical techniques for bulk chemical analysis of materials in several fields (Fitton, 1997). An XRF spectrometer uses primary radiation from an X-ray tube to excite secondary X-ray emissions from the sample. The radiation emerging from the sample includes the characteristic X-ray peaks of major and trace elements present in the sample. Samples were prepared by mixing with a binder, then pressing into pressed powder pellets. Usually the technical apparatus and standards used for major oxides are not the same for trace elements, so that XRF analysis produces two different compositional data sets for the same powdered sample. We applied CODA-CCO to the XRF data set in order to investigate the relationships between the major oxide compositions (%) and the trace element compositions (ppm). Before doing any biplot interpretation, we first comment the numerical output of the analysis given in Table 2. This validation is important, because patterns detected in a biplot are unreliable when the overall goodness-of-fit is low, or if the involved variables are poorly represented. The full space of the solution of this data set has 9 dimensions, and Table 2 provides the numerical output for the first three dimensions of the analysis. The first three canonical correlations are high. This means that the two measurement domains, centred log-ratios of oxides and of trace elements, share variation to a large extent. Statistical significance of the canonical correlations was assessed by means of a permutation test. Such a test is performed by keeping one matrix fixed, say **X**, and randomly permuting the rows of **Y**. The permuted data set is analysed by CODA-CCO, and the canonical correlations are registered. This procedure is repeated 10,000 times and in this way the distribution of the canonical correlations under the null hypothesis of no association between *X* and *Y* is generated. The observed canonical correlations of the original data set are compared against the generated distribution, and a p-value is calculated as the percentage of times the generated values exceed the observed canonical correlations. We found all nine canonical correlations to be highly significant with vanishingly small p-values. Results of the permutation test are given for all nine dimensions in supplementary Figure S2. Test results suggest all nine dimensions potentially could have a geological interpretation, though for reasons of space we limit ourselves to interpreting the first two dimensions.

**Table 2:** Canonical correlations (*r*_1_, *r*_2_, *r*_3_), canonical weights, canonical loadings (between parentheses), adequacy coefficients (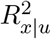, and cumulative 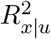) and redundancy coefficients (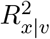, and cumulative 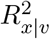) obtained in a CODA-CCO of major oxides and trace elements of European floodplain sediments. Major oxides and trace elements with a canonical coefficient larger than 0.25 in absolute value are marked in bold for the first two dimensions.

| | $r_1 = 0.936$ | | $r_2 = 0.895$ | | $r_3 = 0.814$ | |
| --- | --- | --- | --- | --- | --- | --- |
| | $U_1$ | | $U_2$ | | $U_3$ | |
| SiO<sub>2</sub> | **-0.253** | **(-0.653)** | 0.092 | (-0.366) | -1.501 | (-0.556) |
| Al<sub>2</sub>O<sub>3</sub> | 0.110 | (-0.056) | 0.092 | (-0.446) | 0.401 | (0.372) |
| Na<sub>2</sub>O | -0.057 | (-0.344) | **0.263** | **(-0.234)** | 0.438 | (0.439) |
| MgO | 0.145 | (0.415) | 0.083 | (0.357) | 0.071 | (0.426) |
| P<sub>2</sub>O<sub>5</sub> | 0.017 | (0.001) | 0.028 | (-0.016) | -0.140 | (-0.328) |
| K<sub>2</sub>O | **-1.524** | **(-0.812)** | -0.145 | (-0.481) | 1.314 | (0.190) |
| CaO | -0.033 | (0.112) | **0.551** | **(0.918)** | -0.096 | (-0.227) |
| TiO<sub>2</sub> | **0.738** | **(0.236)** | **-1.069** | **(-0.838)** | -0.840 | (-0.094) |
| MnO | -0.054 | (0.302) | -0.244 | (-0.404) | -0.180 | (-0.263) |
| Fe<sub>2</sub>O<sub>3</sub> | **0.910** | **(0.628)** | 0.350 | (-0.488) | 0.532 | (-0.019) |
| $R_{x\vert u}^2$ | 0.193 | | 0.269 | | 0.110 | |
| $R_{x\vert u}^2$ | 0.193 | | 0.463 | | 0.572 | |
| $R_{x\vert v}^2$ | 0.169 | | 0.216 | | 0.073 | |
| $R_{x\vert v}^2$ | 0.169 | | 0.385 | | 0.458 | |
| V | **0.579** | **(0.693)** | -0.114 | (-0.053) | 0.426 | (0.334) |
| Cr | -0.067 | (0.467) | 0.106 | (0.191) | -0.144 | (-0.182) |
| Co | **0.372** | **(0.700)** | **-0.399** | **(0.032)** | 0.180 | (0.154) |
| Ni | 0.012 | (0.577) | **0.266** | **(0.324)** | -0.215 | (-0.042) |
| Cu | -0.046 | (0.450) | 0.074 | (0.284) | -0.125 | (-0.158) |
| Zn | 0.144 | (0.281) | -0.006 | (-0.016) | -0.239 | (-0.289) |
| Ga | **0.541** | **(-0.110)** | -0.214 | (-0.104) | 0.858 | (0.670) |
| As | 0.091 | (0.181) | 0.038 | (-0.201) | -0.133 | (-0.231) |
| Rb | **-1.848** | **(-0.762)** | 0.138 | (-0.189) | 1.309 | (0.517) |
| Sr | -0.137 | (-0.153) | **1.200** | **(0.859)** | -0.087 | (0.108) |
| Zr | **-0.443** | **(-0.463)** | -0.218 | (-0.292) | -1.236 | (-0.441) |
| Nb | **0.622** | **(-0.046)** | **-0.544** | **(-0.284)** | -0.220 | (0.102) |
| Sn | 0.089 | (-0.046) | -0.067 | (-0.091) | -0.011 | (-0.012) |
| Cs | -0.057 | (-0.502) | -0.107 | (-0.151) | -0.303 | (-0.195) |
| Ba | -0.140 | (-0.507) | **-0.450** | **(-0.265)** | 0.105 | (0.162) |
| Pb | -0.097 | (-0.168) | 0.114 | (-0.109) | -0.017 | (-0.307) |
| Th | **0.401** | **(-0.410)** | 0.191 | (-0.222) | -0.275 | (0.208) |
| U | -0.016 | (-0.351) | -0.007 | (-0.141) | 0.126 | (0.342) |
| $R_{y\vert v}^2$ | 0.194 | | 0.078 | | 0.088 | |
| $R_{y\vert v}^2$ | 0.194 | | 0.272 | | 0.360 | |
| $R_{y\vert u}^2$ | 0.170 | | 0.062 | | 0.058 | |
| $R_{y\vert u}^2$ | 0.170 | | 0.233 | | 0.291 | |

Inspection of Table 2 shows that the oxides SiO_2_, K_2_O, Fe_2_O_3_ and TiO_2_ and the trace elements V, Co, Rb, Ga, Zr, Nb and Th are important contributors to the first dimension of the analysis. For the second dimension these are the oxides CaO and TiO_2_, trace Sr, and to a lesser extent trace elements Zr, Ni, Nb, Ba, Co and Ga. We focus on parts with canonical coefficient above 0.25 in absolute value. Often, though not always, these parts also have large canonical loadings. When interpreting the biplot, we will mainly focus on links involving these components. The biplot of the analysis is shown in Figures 4A (major oxides in standard coordinates) and 4B (trace elements in principal coordinates). Plots 4A and 4B can be overlaid, but are presented separately to avoid an overcrowded display. We again look for links that run parallel to the canonical variables, which are represented by the perpendicular coordinate axes. Such links, representing approximately the standard deviation of the logarithm of ratios, can be expected to have particular strong correlations. The first three canonical correlations are 0.94, 0.89 and 0.81. Numerical output of the CODA-CCO indicates that the two dimensional biplot accounts for 44% of the variation in the between-set covariance matrix of the clr transformed compositions, accounting, in the scaling used, for 46% of the total variance of clr transformed oxides, and 23% of the total variance of clr transformed trace elements. Log-ratios of the oxides that have large correlation with the first canonical variate are ln(K_2_O/Fe_2_O_3_), ln(K_2_O/TiO_2_), and ln(SiO_2_/Fe_2_O_3_), while for the trace elements these are ln(Rb/V), ln(Rb/Co), and ln(Ba/Co). The association among K_2_O, SiO_2_, Rb and Ba for negative values of the first canonical variate has a well defined geochemical meaning. They trace the behavior of lithofile elements that follow Potassium geochemistry and the relative increase in Silica content (mainly presence of K-Feldspars) in the bedrock nature across Europe. On the other hand, positive values associated with Fe_2_O_3_, TiO_2_, V and Co point out the presence of mafic and ultramafic lithologies (relative decreasing Silica content) as well as mineralizations and presence of clay-rich soil with relatively high Al_2_O_3_ contents. The second canonical variate points out the association among Ca and Sr versus that of TiO_2_, Fe_2_O_3_, MnO, Nb, Ga and Ba. This shows the presence of carbonatic lithologies versus the presence of mafic and ultramafic rocks, felsic crystalline rocks or clay-rich soils with high Al_2_O_3_ contents, as well as the presence of mineralization (i.e. Ba). We confirmed the relationships between log-ratios inferred from the biplot by making a scatterplot matrix, where the most prominent log-ratios are shown (see supplementary Figure S3 for log-ratios related to the first canonical variate).

**Figure 4:**
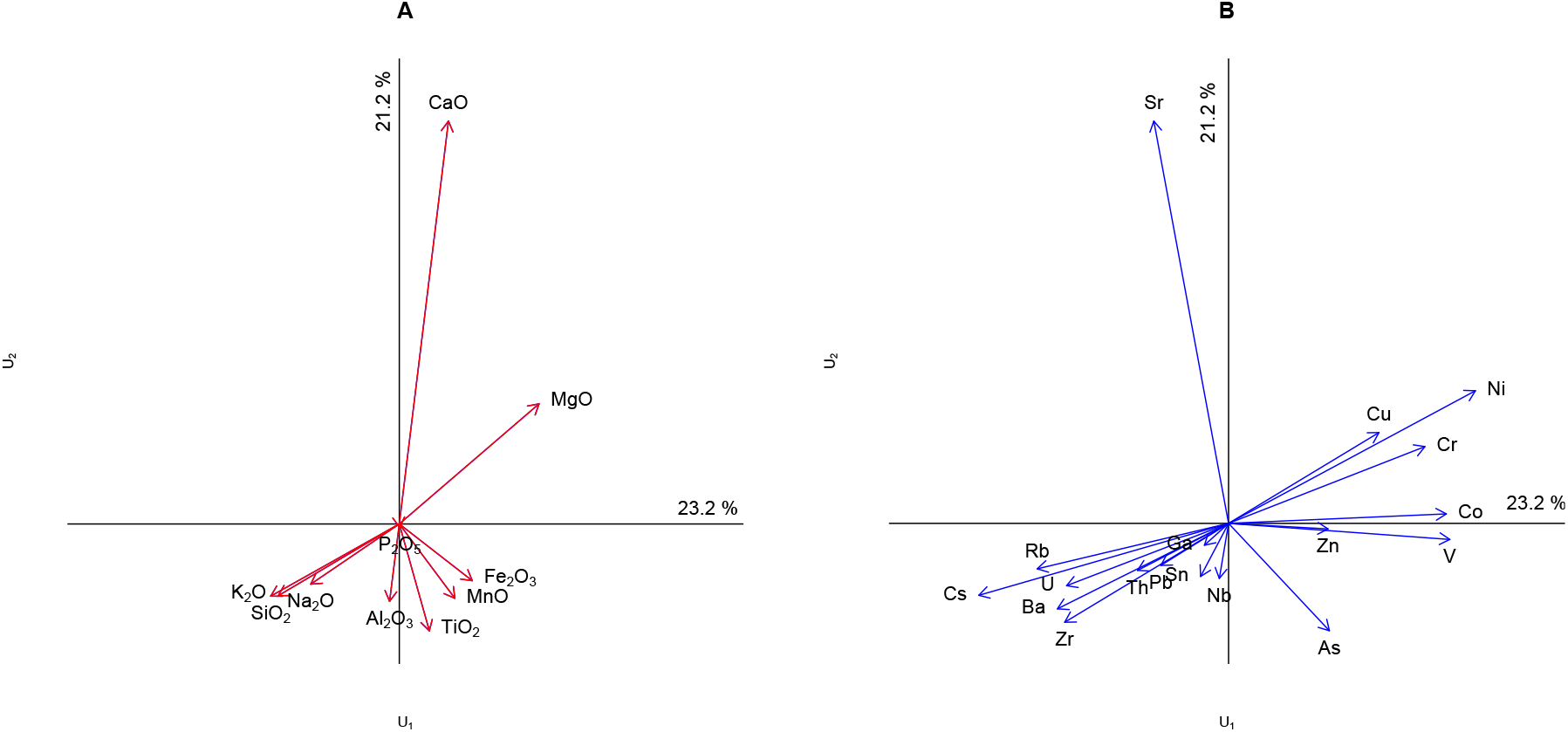
CODA-CCO biplot of major oxides (in standard coordinates) and trace elements (in principal coordinates). Rays represent clr-transformed parts.

When the canonical correlation analysis is stratified on a country-wise basis (biplots and tables not reported), using countries where a sufficient numbers of samples is available, interesting features emerge that can be related to general geochemical laws. In fact, the major oxides K_2_O and Rb are closely related to the first canonical variate across all countries (ignoring Austria and Poland because of small sample size), while Al_2_O_3_ and Sr are quite related to the second one. The result describes well the geochemical affinity between K_2_O and Rb. Both the elements pertaining to group 1 of the periodic table and the Rb+ ion (ionic radius 152 pm) substitute for K^+^ (138 pm) in several minerals, thus tracing its geochemical distribution in outcrops across all Europe. The association between Al_2_O_3_ and Sr appears to point out sedimentary processes, where the distribution of Sr may be affected by strong adsorption on clay minerals containing Al_2_O_3_. The 747 samples can be projected onto the biplot in Figure 4 and this can aid interpretation (Graffelman, 2005). Supplementary Figure S4 shows this more dense biplot. This plot shows some French samples (top-right) which are relatively high in Sr, MgO and CaO, a set of Spanish samples relatively high in Sr, and a set of Polish samples which are relatively low on the first canonical variate. However, in general, the samples of the different countries overlap to a large extent.

CODA-CCO biplots based on a pairwise log-ratio approach are shown in Figure 5, where oxides (5A and 5C, in standard coordinates) and traces (5B and 5D, in principal coordinates) are presented separately. Figures 5A and 5B show the covariance-based analysis, whereas Figures 5C and 5D show the correlation-based analysis. 5A and 5B should be overlaid for interpretation, and 5C and 5D too. We emphasize that between-set scalar products of Figures 5A and 5B approximate between-set covariances, whereas between-set scalar products of Figures 5C and 5D approximate correlations. Because of the large canonical correlations, there is little difference between the use of standard and principal coordinates. Because there are so many pairwise log-ratios, the pairwise log-ratios in Figure 5 were filtered by goodness-of-fit, and only those log-ratios that have 75% or more of their variance accounted for are shown. For log-ratios in principal coordinates, this threshold was lowered to 60%, as the goodness-of-fit in this scaling is typically worse. We summarize the main relationships uncovered by these biplots: ln(K_2_O/TiO_2_) and ln(K_2_O/Fe_2_O_3_) are positively correlated, and have strong negative correlation with two log-ratios involving Rb, ln(Co/Rb) and ln(V/Rb). Samples with high values on the latter two log-ratios have low values on the log-ratios ln(K_2_O/TiO_2_) and ln(K_2_O/Fe_2_O_3_). This is the most salient feature of the dataset uncovered by the first canonical variate. These relationships were also uncovered in the previous clr-based analysis. The second canonical variable is associated with at least six log-ratios that all involve CaO, and at the same time with at least eight traces that all involve Sr. The biplots in Figure 5 represent in fact two approximately orthogonal sets, if CaO and Sr are consistently placed in the numerators (or denominators) of all involved log-ratios. Note that the results are consistent with the clr-based analysis, where CaO and Sr had the longest biplot vectors and therefore many long links involving these components. The goodness-of-fit of the between-set covariance and correlation matrices in Figure 5 is 44.4%, and coincides with the goodness-of-fit of the between-set clr covariances in Figure 4.

**Figure 5:**
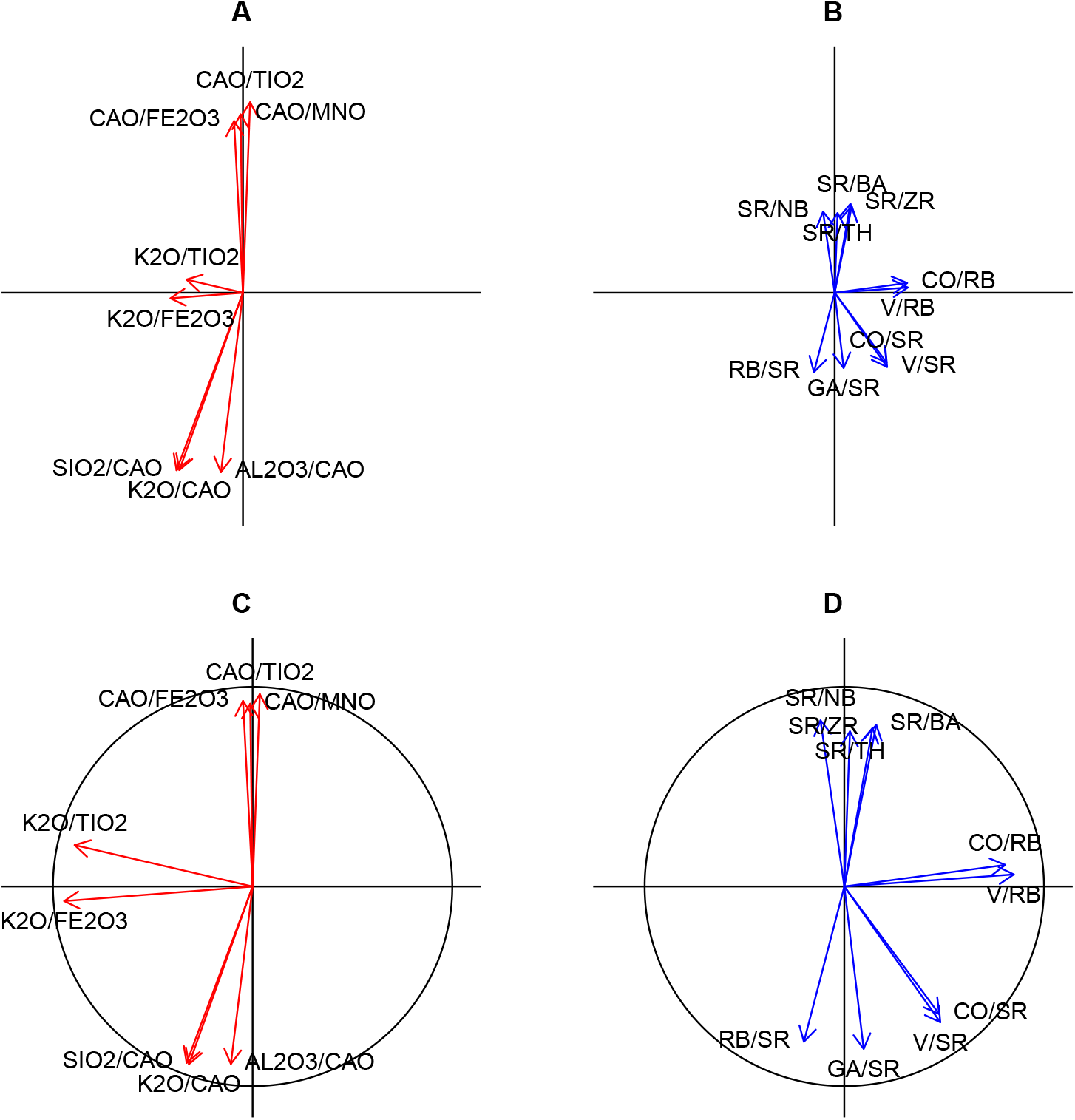
CODA-CCO biplot of major oxides (in standard coordinates) and trace elements (in principal coordinates). Rays represent pairwise log-ratios. Panels A and B show a covariance-based analysis, whereas C and D show a correlation-based analysis.

## 4 Conclusions and discussion

Compositional canonical correlation analysis (CODA-CCO) has been presented as a technique for analysing two compositional datasets, an *X* set and a *Y* set, potentially measuring different kinds of compositions, e.g. *X* may refer to a microbial composition and *Y* to the biochemical composition of the same set of subjects. We note that the method presented here has a wider scope of application, because it can also be applied to two subcompositions (that do not share parts) build from the *same* composition. The method will then act as a magnifying glass focusing on the relationships between the two subcompositions.

Canonical correlation analysis provides the best approximation, in the generalized least squares sense, to the between-set covariance matrix. In the context of compositional data, as treated in this paper, the CODA-CCO biplot gives the optimal approximation for the between-set covariance matrix of the clr transformed parts in the clr-based approach, and the optimal approximation for the between-set covariance matrix of the pairwise log-ratios in the plr based approach. The same goodness-of-fit is obtained for both covariance matrices, as justified by Equation (24), and as could be observed in both examples. Biplots in both approaches are fully equivalent if data is not standardized. If all within-set links in the clr-based biplot are “extracted” by calculation of all possible difference vectors, and plotted as vectors emanating from the origin, then the biplot of the pairwise log-ratio approach will be obtained. This property holds for any chosen dimensionality. Because the representation of the links is explicitly optimized in the pairwise approach, it follows that the links are also optimally displayed in the clr-based biplots. This equivalence is clearly visible by comparing the full space solutions in Figures 2A and 2B with Figures 3A and 3B, but it also holds for approximate solutions like the ones given in Figures 4A, 4B and 5A, 5B respectively.

The aforementioned equivalence between the clr and plr biplot may suggest the latter to be superfluous, but in our opinion this is not the case for several reasons. First, the clr-based approach is limited in the sense that links always represent pairwise log-ratios. In the plr approach, links between pairwise log-ratios correspond to balances, and the pairwise approach may uncover the existence of balances, or even correlations between balances, so allowing for a richer and more refined analysis. If the analysis is limited to the clr biplot, potentially interesting balances that invoke more than two parts may go unnoticed. Second, in the clr-based approach correlations between log-ratios are not optimally displayed, whereas they can be optimally represented in the pairwise approach. By standardization of the pairwise log-ratios by division by their standard deviation prior to canonical analysis, biplots with an optimal approximation to the correlation structure of the pairwise log-ratios are obtained. In the latter plots, unit circles are illustrative, as the goodness-of-fit of the pairwise log-ratios can be inferred from the ray’s length. We note that these considerations carry over to compositional biplots made by principal component analysis as well.

Biplots are not unique, and in practical data analysis it may be daunting to choose the most appropriate plot for representing a given data set. This is particularly true for the rich family of compositional canonical biplots proposed in this paper. The analyst is confronted with at least three decisions in the analysis: a) whether to use a clr or plr based approach, b) to standardize the data prior to analysis or not and c) whether to use standard coordinates for rows and principal coordinates for columns or the other way around. We present some considerations on these issues, hoping this will help analysts to make a sensible choice.

The clr-based approach has the advantage of producing less dense plots having fewer rays. In principle, all pairwise log-ratios are present in this biplot by means of the links. The analyst will have to make the mental effort to search for interesting links, in particular by looking at links that run parallel to canonical variables with a high correlation. Standardization by division by the standard deviation of the clr transformed data, prior to the canonical analysis will differentially scale the columns of the clr transformed data. This complicates the interpretation of the links, and seems therefore generally not indicated. Regarding the biplot scaling, if there is particular interest in representing the within-set correlation of one set, that set should be represented in standard coordinates to enhance the representation of its correlation structure. If both sets are equally important, then a pragmatic rule is to choose that scaling that explains most of the total variance of the data, as expressed by the adequacy and redundancy coefficients. For instance, biplot 2A of the artificial data in Section 3.1 explains 100% and 36.3% of the variance of the *X* and *Y* data in F_s_G_p_^;^ scaling, but respectively 81.5% and 100% in F_p_G_s_^;^ scaling. The latter may be preferred for giving, overall, a better approximation to the transformed data matrices. To safeguard against erroneous interpretations, we recommend always to explore the data using both biplot scalings used in this paper (Equations (3) and (4)). Patterns like parallel links that show up in *both* biplots are more likely to be real.

The pairwise log-ratio biplot has the advantage that pairwise log-ratios are directly displayed as rays in the biplot. With large compositions the number of links can be prohibitive, and produce very dense biplots. However, as shown in the geological example in subsection 3.2, by removing all links with a low goodness-of-fit these plots can be improved, and salient features of the data can be made visible.

In compositional data analysis, several log-ratio transformations are in use. In particular, the isometric log-ratio (ilr) transformation is increasing in popularity, as it provides Cartesian coordinates to represent the compositions. We show in Appendix A that a CODA-CCO of the ilr transformed compositions will yield the same canonical correlations and the same canonical variables. An advantage of using ilr coordinates is that the singularity of the covariance matrices is avoided, which frees one from the need to calculate generalized inverses. For biplot construction however, a clr based or plr based analysis seems to be the most straightforward approach to CODA-CCO. Given that the clr, plr and ilr transformations are linearly related to each other, the proposed compositional biplots could also be derived from a canonical analysis of ilr transformed compositions.

## Acknowledgments

This work was supported by grants MTM2015-65016-C2-1-R and MTM2015-65016-C2-2-R (MINECO/FEDER) of the Spanish Ministry of Economy and Competitiveness and European Regional Development Fund, by grant 2030_M1488-BUCBASI09 of PROG-GEOBASI TOSCANA, by the University of Florence funds, 2015 and 2016, and by Grant GM075091 from the United States National Institutes of Health.

## Appendices

## A Appendix

In this appendix we show the invariance of the canonical correlations and the canonical variables when the ilr transformation is used instead of the clr transformation. The singular value decomposition in Equation (22) can be rewritten as an equivalent eigenvalue decomposition:

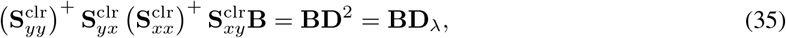

where D_λ_ contains the eigenvalues (squares of the singular values) of the spectral decomposition. The clr and ilr coordinates are linearly related by the following expressions (Egozcue et al., 2003)

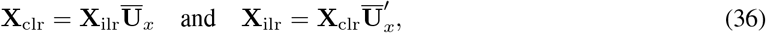

where 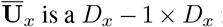 is a *D_x_* — 1 × *D_x_* matrix with orthonormal rows, satisfying 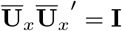 and 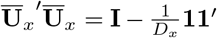. We use the 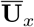 notation in order to follow the usual notation in CODA (Egozcue et al., 2003), but put a bar in order not to create confusion with the previously defined canonical *X* variables (**U**), and use a subindex *x* to show that it applies to the *X* composition. Note that 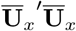 is an idempotent centring matrix. The analogous transformation for the *Y* variables is given by a *D_y_* — 1 × *D_y_* matrix 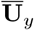, as the *X* and *Y* set may not have the same number of parts. By substitution we obtain straightforward expressions for the relationships between ilr and clr within-set and between-set covariance matrices of the *X* and *Y* compositions, given by:

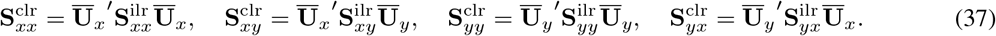

Premultiplication by 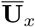 and postmultiplication by 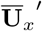 (or 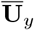 and 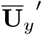, as corresponds) allows us to obtain the ilr covariance matrices from the clr covariance matrices:

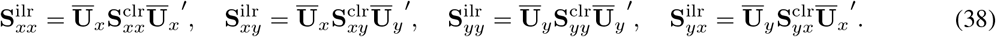

Substituting (37) in (35) gives

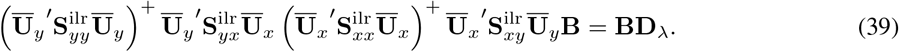

At this point we note that 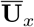 has rank *D* — 1, and that the rows of 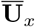 are linearly independent. In that case, the Moore-Penrose inverse of 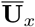 is given by

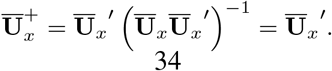

Similarly, we also have 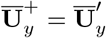. Equation (39) can now be simplified to

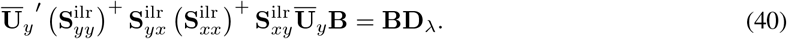

Since the covariance matrices of the ilr coordinates are invertible, premultiplying by 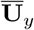 this can be rewritten as

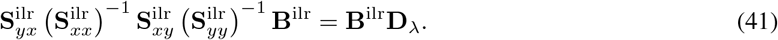

with 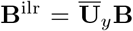, satisfying 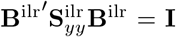. Equation (40) is the eigenvalue-eigenvector decomposition corresponding to a canonical correlation analysis of *X* and *Y* compositions in ilr coordinates. Finally, canonical variables obtained in the clr based and in the ilr based approach will be identical because 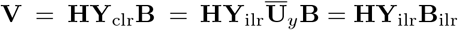. Equation (40) shows that a clr-based and ilr-based CODA-CCO yield the same canonical correlations, yield canonical coefficients that are related by a linear transformation, and yield the same canonical variables. Equations (40) and (41) show that in the clr-based approach the canonical coefficients of one canonical variate (columns of matrices A and B) sum to zero. Since 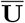 satisfies 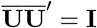 and 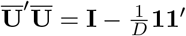 we have that 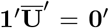. From Equation (40) it follows that the columns of **B** sum to zero. By using a spectral decomposition analogous to Equation(35) with **A** as eigenvectors, the same property can also be shown for **A**.

## Supplementary figures

**Scatterplot matrix of pairwise log-ratios of artificial 3-part compositions**

**Figure S1:**
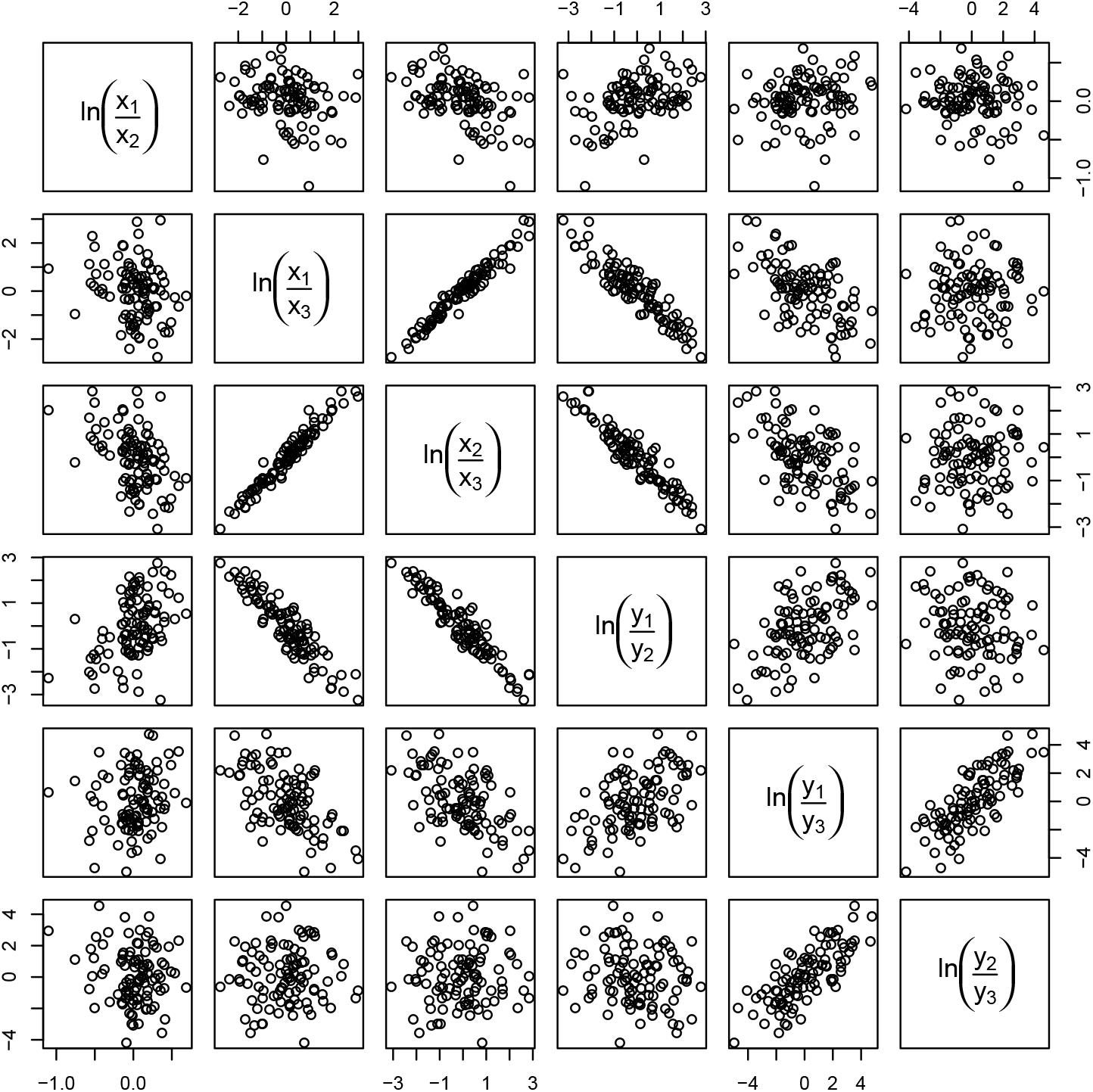
Scatterplot matrix of all log-ratios of two compositions of three parts.

**Results of permutation test for European floodplain sediments**

**Figure S2:**
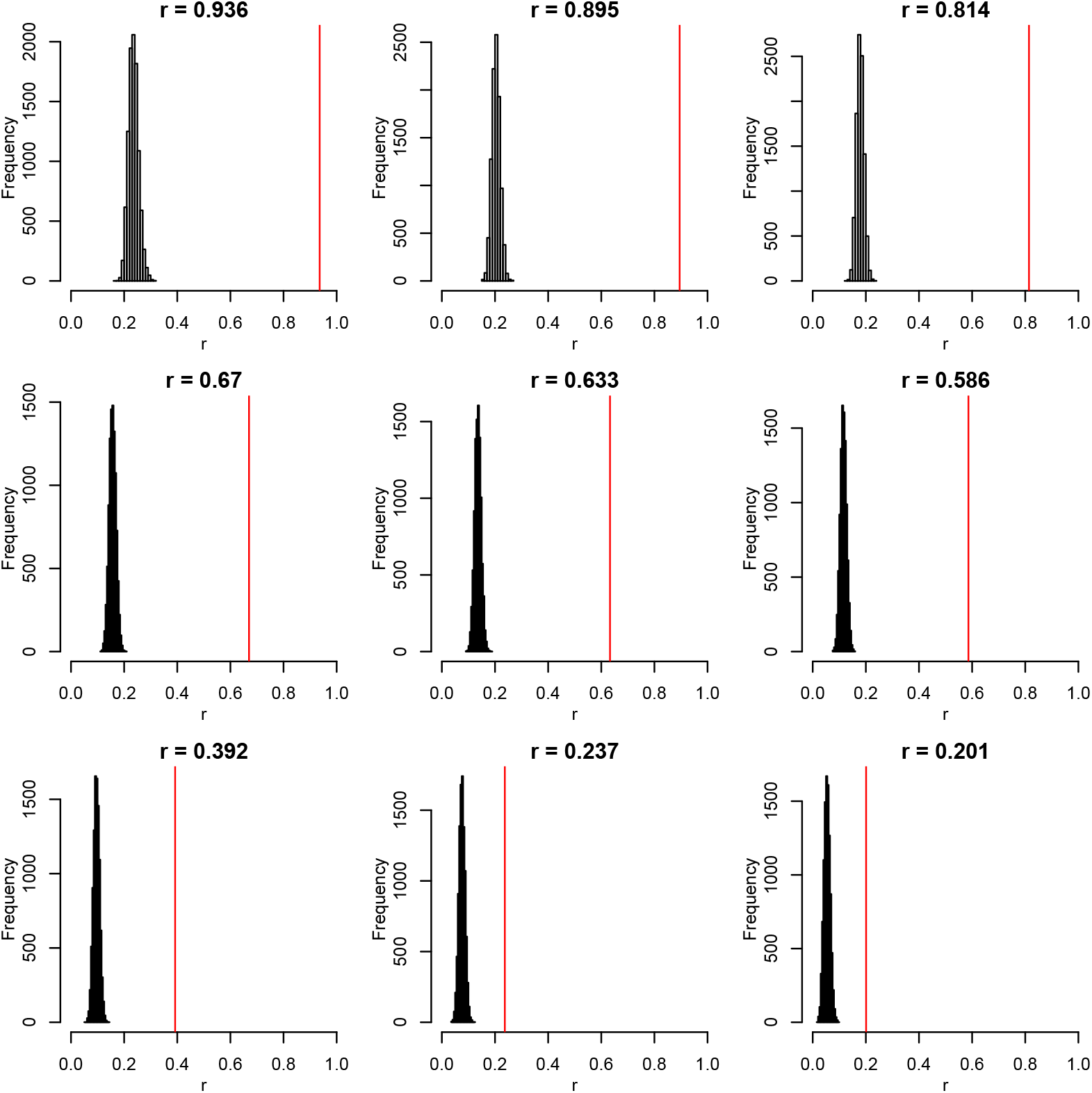
Permutation test of nine canonical correlations obtained in a CODA-CCO of oxides and trace elements of European floodplain sediments. Histograms show the distribution of the canonical correlation obtained in 10,000 permutations of the data. The vertical red line indicates the observed sample canonical correlation.

**Scatterplot matrix of selected pairwise log-ratios for European floodplain sediments.**

**Figure S3:**
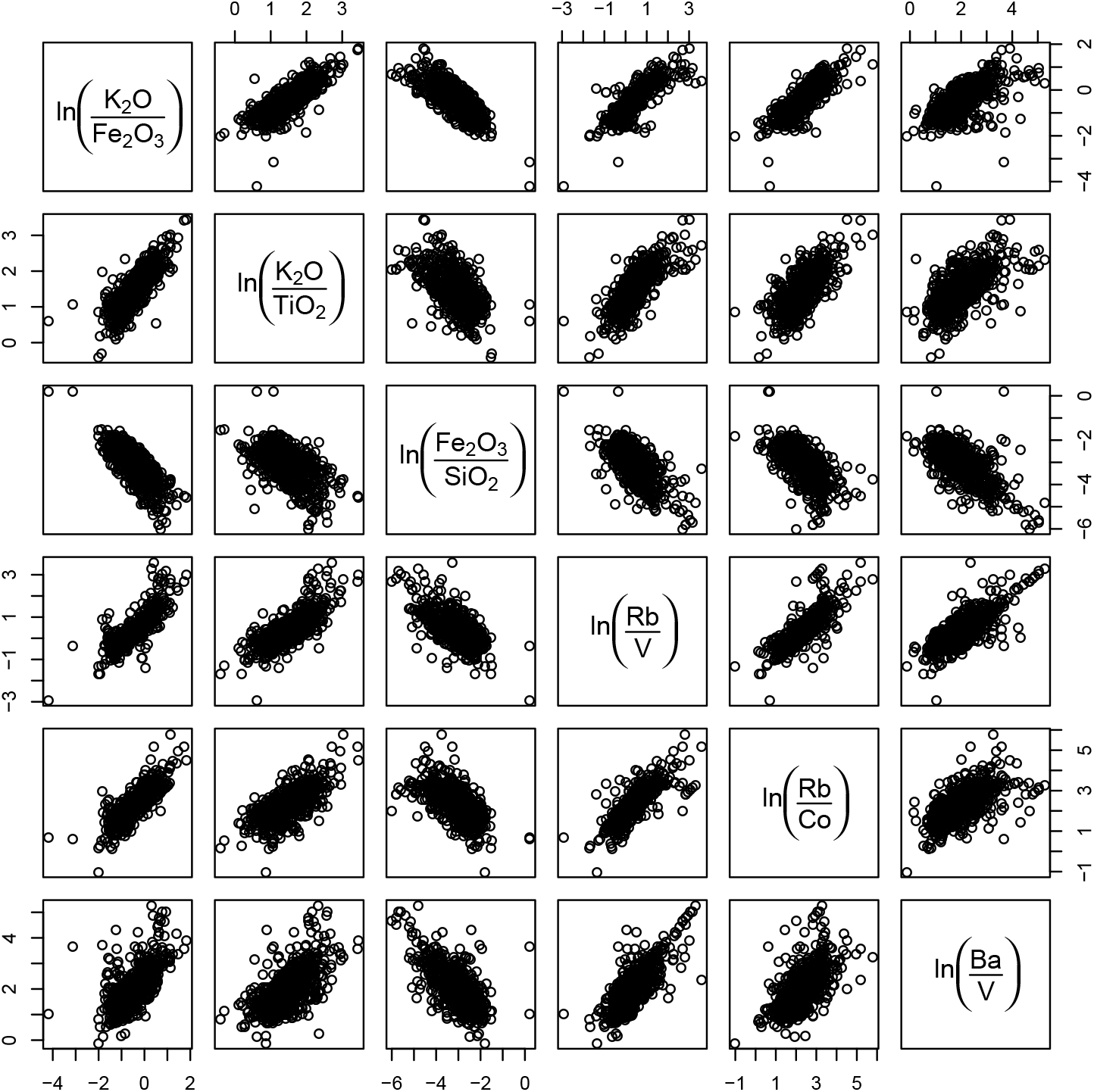
Scatterplot matrix of log-ratios of major oxides and trace elements of European floodplain sediments related to the first canonical variate.

**Biplot enhanced with samples for European floodplain sediments.**

**Figure S4:**
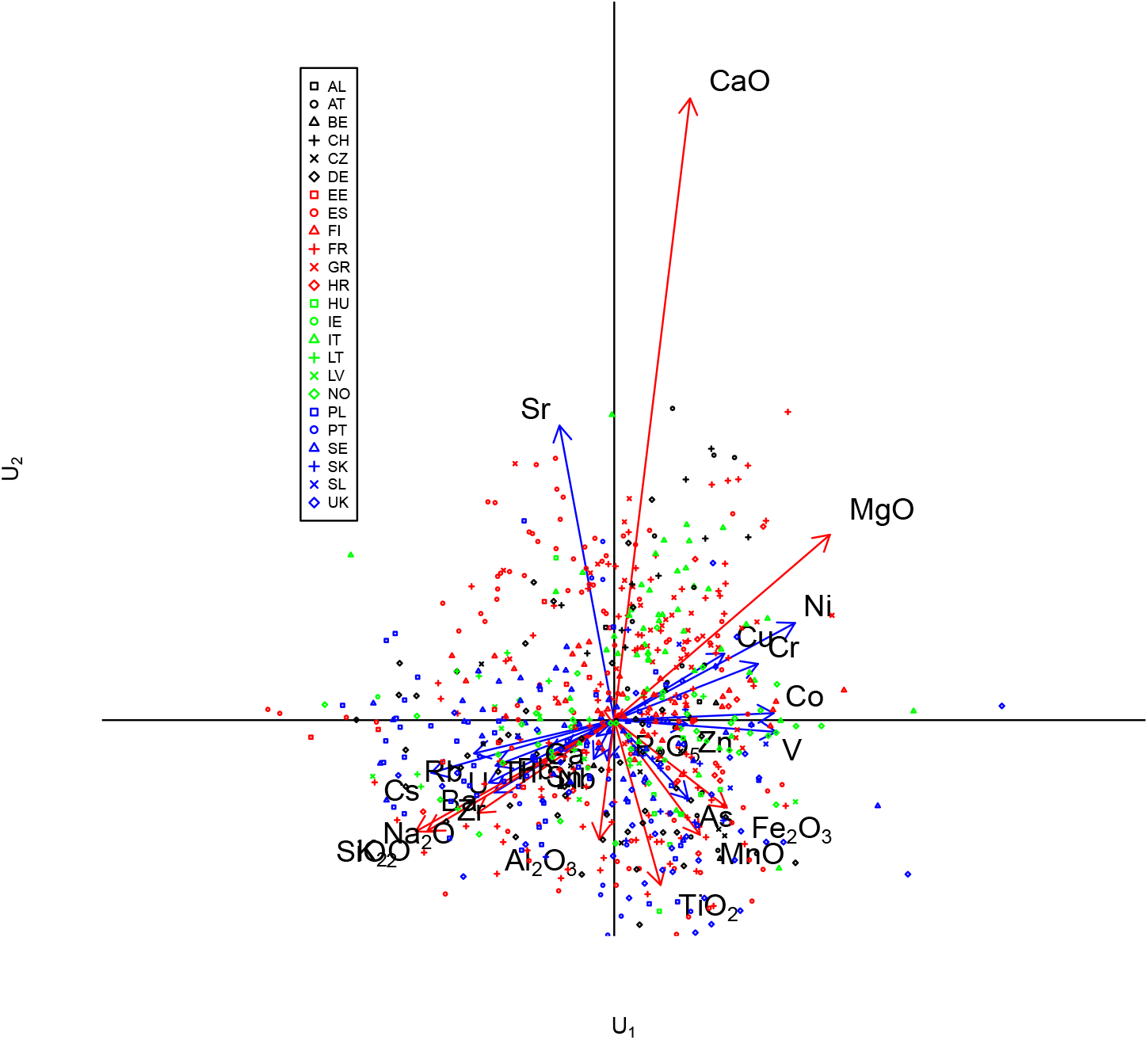
Biplot of CODA-CCO of major oxides and trace elements of European floodplain sediments, with added samples. Symbols and colour are used to indicate country of origin.

